# Uterine secretome initiates growth of gynecologic tissues in ectopic locations

**DOI:** 10.1101/2023.10.04.560850

**Authors:** Jan Sunde, Morgan Wasickanin, Tiffany A Katz, Sanam Bidadi, Derek O’Neil, Richard O Burney, Kathleen A Pennington

## Abstract

Endosalpingiosis (ES) and endometriosis (EM) refer to the growth of tubal and endometrial epithelium respectively, outside of their site of origin. We hypothesize that uterine secretome factors drive ectopic growth. To test this, we developed a mouse model of ES and EM using tdTomato (tdT) transgenic fluorescent mice as donors. To block implantation factors, progesterone knockout (PKO) tdT mice were created. Post-ovulatory endometrium was induced hormonally in donor tdT and wild-type (WT) female mice. tdT oviductal cells and WT endometrium were harvested for intraperitoneal injection into synchronized recipient WT mice. Ectopic lesions were identified using fluorescence in-vivo imaging and then harvested for histological evaluation. Fluorescent lesions were present after oviduct implantation with and without WT endometrium. Implantation was increased (p<0.05) when tdt oviductal tissue was implanted with endometrium compared to oviductal tissue alone. Implantation was reduced (p<0.0005) in animals implanted with minced tdT oviductal tissue with PKO tdT endometrium compared to WT endometrium. In conclusion, endometrial derived implantation factors are necessary to initiate ectopic tissue growth. We have developed an animal model of ectopic growth of gynecologic tissues in a WT mouse which will potentially allow for development of new prevention and treatment modalities.

## Introduction

Endosalpingiosis (ES), the growth of uterine tubal tissue ectopically, has been an area of intense study recently following a 2016 retrospective chart review found ES associated with gynecological malignancy in 42% of patients, with a prevalence of approximately 1.5% in 60,000 gynecologic specimens [1]. With intensive pathologic evaluation, we subsequently reported an increased premenopausal prevalence of benign ectopic growths, including ES (37%), endometriosis (EM) (32%), paratubal cysts (47%), and Walthard’s nests (ectopic urothelial cell growths) (29%) in women with at-risk tissues age 31-50, with ES prevalence increasing to 66% after menopause, in contrast to a decrease in EM (5%), while 89% of post-menopausal specimens had some type of ectopic lesion and various lesion types could be present in the same patient [2]. Another recent study reporting EM in 39% of random biopsies of patients with chronic pelvic pain also found much higher prevalence than expected [3].Sampling bias is the likely explanation for retrospective studies reporting an association of these benign peritoneal cavity cellular implants with malignancy [4–6], since the large majority of patients undergoing surgery for benign conditions do not have a histologic evaluation of all tissues as thorough as that performed for malignancy.

A leading theory for EM development is retrograde menstruation, where during menstruation endometrial cells travel via the fallopian tubes into the pelvic cavity via retrograde flow. However, retrograde menstruation occurs in up to 90% of women, while historically only 6-10% of reproductive age women were reported to develop EM [7]. Other factors at play in the development of EM may depend on factors present in the peritoneal fluid, likely migrating from the uterus. A multitude of studies have identified increased levels of a variety of cytokines and growth factors, including, interleukin-1β (IL-1β), IL-6, IL-8, tumor necrosis factor-α (TNF-α), epidermal growth factor (EGF), Fibroblast Growth Factors (FGFs), vascular endothelial growth factor (VEGF), and leukemia inhibitory factor (LIF) in peritoneal fluid of women with EM [8–11]. Interestingly, many of these factors are essential to the process of implantation, including LIF [12].

Therefore, upon considering the ubiquity of benign ectopic growths and their coincidental association with malignant conditions, in addition to the retrograde menstruation theory of EM development [7], the unexplained facts that bilateral tubal ligation and also hysterectomy decrease the risk of ovarian cancer [13–16], and that uterine secretome factors are present in peritoneal fluid of women with ectopic lesions [8–10] we hypothesized that cyclic post-ovulatory uterine secretome factors in the uterine fluid drive free-floating epithelial cells to implant and grow in ectopic locations.

To test this hypothesis, animal models are needed. Normal mouse strains and genetically mutated models of ovarian inclusion cysts (OIC, ES of the ovary) mimic findings we and others report in humans, such as increasing ES incidence with age [4, 17–19]. Specified genes have been demonstrated to play a role in the pathogenesis of ovarian inclusion cysts in transgenic models, but these models may also unexpectedly alter genes normally involved in lesion development and these models are not designed to evaluate the role of uterine secretome factors in ectopic lesion development. Furthermore, while some mouse models of ectopic growth (EM models) have begun to incorporate fluorescent mouse strains to allow for in-vivo visualization of ectopic lesion growth, there is a need to optimize these models in order to track lesion progression over time [20]. Therefore, the goals of our current study were two-fold, (1) to refine current mouse models of ectopic tissues to better visualize in-vivo lesion initiation and (2) determine the effects of uterine secretome factors on ectopic lesion growth in our mouse model. To do this we developed a transgenic fluorescent mouse model using tdTomato mouse tissues injected into the peritoneum of WT recipient mice to investigate this “uterine secretome” hypothesis, with the hope of identifying key factors involved in the initial step in the pathogenesis of ectopic growths.

## Materials and methods

### Animals

This study was approved by the Institutional Animal Care and Use Committee (IACUC) at the Madigan Army Medical Center (MAMC) and Baylor College of Medicine (BCM). Care and procedures for the mice were in accordance with the Helsinki Declaration and Institutional Guide for Laboratory Animals. Mice were housed under a 12/12-hour light/dark cycle at 25 C and 50% humidity and were fed ad libitum with a standard diet and water. Mature 8-12 week old female mice expressing ubiquitous tdTomato (tdT) from the Ai14flox allele (B6.Cg-*Gt(ROSA)26Sortm14(CAG-tdTomato)Hze*/J #:007914, Jackson Labs, Sacramento, CA) were chosen as tissue donors due to their fluorescence and ease of in-vivo imaging tdT mice [18–20]. Mature 8-12 week old WT C56BL/6J mice from Jackson Laboratory or the BCM mouse facility were used for donor tissue and as recipients to minimize genetic off-target effects.

### Experimental design

To easily identify ectopic lesions in live mice we developed a mouse model using tdT fluorescent mice as donors (Fig. 1) and previously published methods from several prior models of endometriosis in mice that artificially induced menstruation in mice using hormone schedules to induce decidualization and endometrial breakdown [21–23]. Once the tdT mouse model was established for optimal lesion visualization, experiments were performed to determine if the uterine secretome plays a role in ectopic lesion development, culminating with experiments evaluating the effect of an exogenous secretome factor, leukemia inhibitory factor (LIF) on ectopic lesion initiation. All experiments were planned with 10-12 mice per arm.

**Fig 1.**
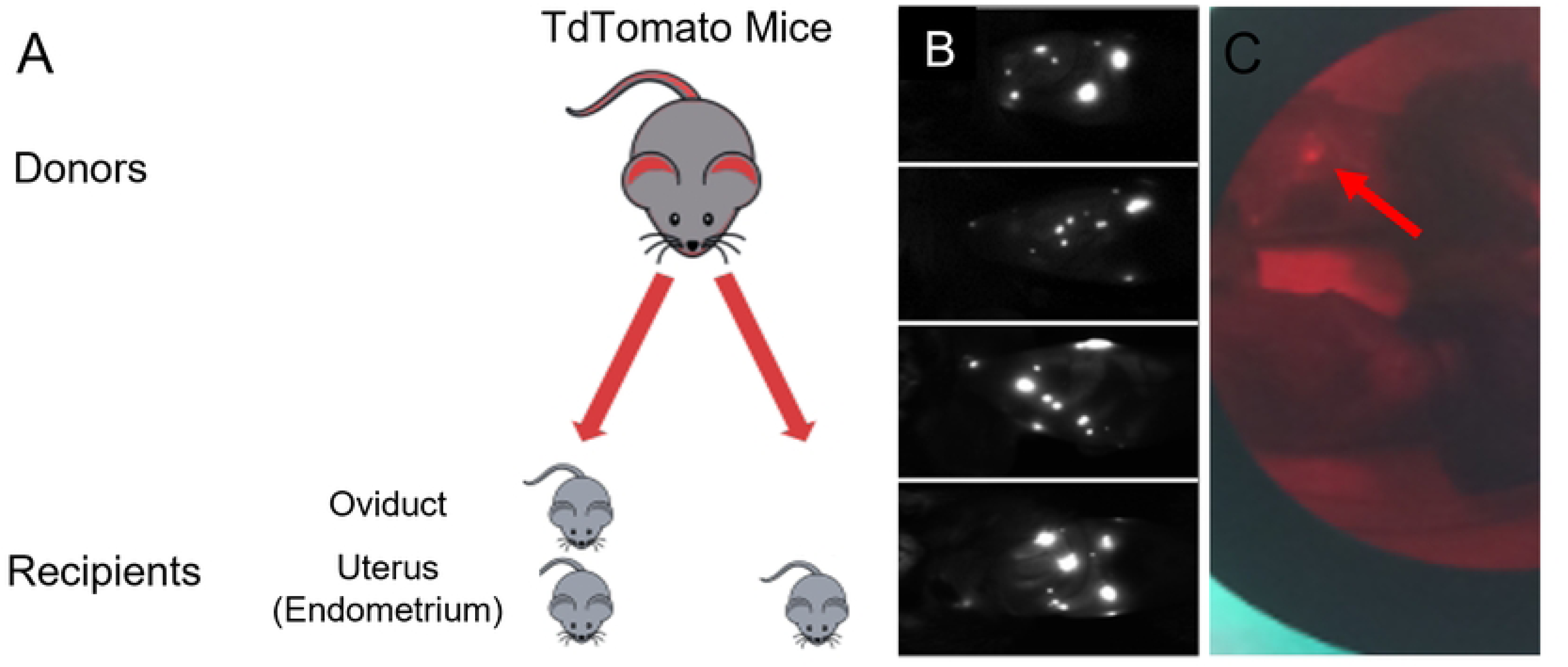
Development of tdTomato mouse model of endometriosis and endosalpingiosis. A. After hormonal synchronization, tdT endometrial and oviductal tissue were collected, processed, and injected into WT recipient mice. B. 28 days post injections recipient mice were imaged with a Kodak in-vivo imaging system, visualized fluorescent light from the implanted endometrial or oviductal tdT tissues representing endometriosis and endosalpingiosis mouse models respectively. C. After euthanization, the fluorescent endometrial or endosalpingeal lesions were counted and collected for histological analysis using direct visualization with green light and red filter.

### Synchronization of donor and recipient animals

Donor animals: Menstrual endometrium and subsequently diestrus (secretory) phase endometrium was induced in tdT and WT C57BL/6J donor mice using a hormonal protocol [24] of daily subcutaneous (SC) injections of estrogen in sesame seed oil (1 ul/g of 100 ng/100 µl via 28 G needle) for three days (days 0-2), followed by SC injections of progesterone (0.5 mg) and estrogen (5 ng) in 100 µl in sesame seed oil daily on days 6-8. On day 8 pseudo-decidualization of the uterine horns was performed under anesthesia by inserting the sesame seed oil into the cervix using a device fashioned from a shortened angiocatheter and a syringe. Daily SC progesterone treatment (0.5 mg) continued for another three days. On day 12, 24-40 hours following the final progesterone injection, mice were euthanized, and tissues were harvested.

Recipient Animals: Female WT C57BL/6 mice were synchronized to diestrus, the cycle phase when secretome factors are expected in the uterine cavity, using progesterone 3ug/200ul saline (15 ug/ml) SC via 28-gauge needle and cloprostenol (prostaglandin F2 alpha), 0.5 ug in 100 ul saline (5 ug/mL) intraperitoneally (IP) via 28-gauge needle followed three days later by a second dose of cloprostenol. This treatment places the recipient mice in the diestrus (post-ovulatory or secretory) phase at the time of tissue injection five days after the initial hormonal injection, which we confirmed histologically. In some studies, male mice were used as recipients in order to eliminate hormonal cycling factors as described below.

### Tissue harvest and implantation

On day 12 of hormone synchronization, donor females were euthanized for harvest of diestrus or menstrual endometrium and/or oviductal and bladder epithelium. Post-mortem, the entire uterine horns, oviducts, and bladder were removed via sterile technique and placed in a petri dish containing warm sterile PBS media. Under a dissecting microscope, each uterine horn was bisected lengthwise, and the endometrium was dissected from the underlying uterine wall. The isolated endometrium was then suspended in warm media. The endometrium was finely minced with scissors as previously described [25]. Oviductal tissues were obtained by dissection and mincing of the oviducts separately in PBS. PBS and tissues were collected in a micro centrifuge tube and centrifuged for 30 – 60 sec at 5000 rpm to concentrate the tissue into a pellet and fluid was removed. Harvesting flushed cells from the oviduct was difficult due to their small size and experimental differences in lesion development were noted using minced tissues, so further experiments were performed with minced tissues to optimize the model. The harvested tissue/cells and/or minced endometrium from donors was suspended in the supernatant and standardized to a volume 400 µl room temperature PBS before intraperitoneal injection into WT recipient mice via a 19-gauge needle. The processing time from the dissection of tissue from donor mice to tissue injection in recipient mice was approximately 20-30 minutes.

It was initially anticipated that the ectopic lesion yield for oviductal tissue would be less than endometrial tissue, planning a 4:1 donor: WT mouse ratio using 2 oviducts/mouse on 2 occasions. In a single early iteration, oviductal cells were isolated by flushing the oviduct. With the assistance of a dissecting microscope, oviducts were trimmed of surrounding membrane and straightened before being placed into a fresh petri dish of PBS. Using an instrument to stabilize one end of the oviduct, a 29-gauge syringe was inserted into the end and flushed with 50 µl of PBS, with collection of the flushed fluid for peritoneal injection. In subsequent experiments, because of the initial extensive lesion development, 1 minced oviduct per recipient mouse (1:2 donor:recipient mouse ratio) was used, with a single peritoneal injection. Minced endometrium from 1 WT diestrus (menstrual phase for initial experiment) uterine horn per recipient mouse was used to provide implantation factors for those mice that were also injected with endometrial tissue (1:2 donor:recipient ratio).

In the initial experiments, donor tissues were injected into the peritoneal cavity of diestrus phase recipient C57BL/6 WT mice, using minced tdT endometrial tissue alone (E), minced tdT oviductal (O) tissue alone, or a combination of tdT oviductal tissue and WT endometrial tissue (EO) from similarly synchronized WT mice donors to provide secretome factors. In all iterations, endometrial, oviductal tissue/cells were harvested from donors and minced/prepared under clean conditions prior to injection into recipient mice. In the third experiment, oviductal cells were recovered by flushing the oviduct. The PBS and oviductal/endometrial tissues were then collected into a micropipette at a volume of 50 µl.

### Optimization and in-vivo evaluation of lesion implantation

WT recipient mice initially underwent live animal (in vivo) imaging under isoflurane anesthesia one week after each minced tissue injection, every 4 weeks with oviductal cells, and prior to post-mortem dissection in the initial experiments to assess for lesion growth prior to harvesting. Critical for the success of the model when using live animal imaging was the optimization of the signal-to-noise ratio in the detection of fluorescent endosalpingiosis lesions. Imaging was performed using a Kodak FX in vivo small animal imaging station (Carestream Health Inc.). Animals were imaged at an excitation wavelength of 555 nm and an emission wavelength of 600 nm for ∼30-120 seconds suited for tdT. Imaging allowed for the visualization of lesions and helped provide insight into the progression of the disease and disease burden initially. Depending on the signal-to-noise resolution and lesion size, it was possible for recipient animals to have negative imaging in the presence of small lesions.

Based on live imaging, animals were euthanized and necropsied on day 28 when implanted with minced tissue, with a delay to day 93 when implanted with oviductal cells to assess for implantation of tdT ectopic lesions. At necropsy, lesions were excised from surrounding tissue with the assistance of fluorescence imaging using the NightSea light (Fig 1C) for counting and histology. The tdT endometrium fluoresces well, allowing for visualization of millimeter size lesions. Fluorescent lesions were counted and harvested post-mortem using a red filter with a 550 nm green light for histologic evaluation by H&E staining. Implantation by imaging and at necropsy was assessed by counting the number of lesions, which could be found anywhere in the peritoneal cavity. To minimize abdominal auto-fluorescence from animal feed, the recipient mice were administered a low-fluorescent, alfalfa free purified diet for 15 days prior to any imaging.

### Histology

All lesions and uterine tissue were dissected and fixed in 10% buffered formalin at room temperature for 24 hours. The tissues were processed through ethanol dilutions into paraffin wax. Transverse 5 µm sections were serially cut. To determine gross histology, sections were stained with hematoxylin and eosin (H&E) as per standard protocols (Fig 2C).

**Fig 2.**
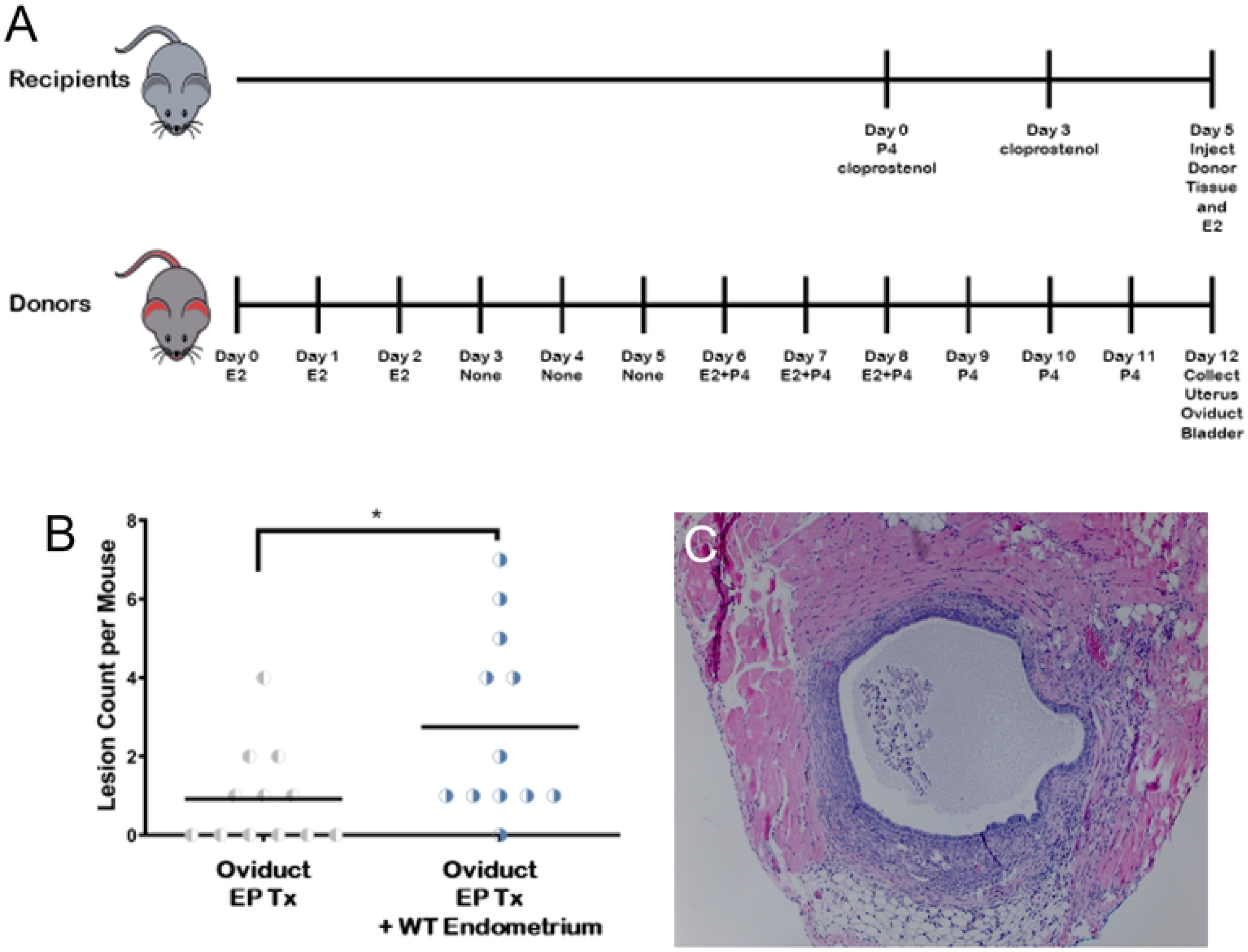
Co-injection of WT endometrium with oviduct increases lesions ectopic lesions. A. Timeline for hormonal induction of menstrual/diestrus endometrium for tdT transgenic donors and WT C57BL/6J female recipients. Induction of menstrual/diestrus endometrium. Donor tdT transgenic female mice were hormonally induced with estrogen and progesterone (EP) to develop menstrual/secretory endometrium. WT C57BL/6J recipients were induced using cloprostenol to develop synchronized diestrus endometrium, using experimental and control groups. Endometrial and oviductal tissue were collected, processed, and injected into recipient mice. E2= estrogen, P4= progesterone B. The number of implanted oviductal lesions in mice had a statistically significant increase with co-injection of WT endometrium in comparison to solely implanted oviductal lesions. C. Histologic analysis confirmed the origin of dissected lesions. Histologic findings using H&E staining of dissected lesions shows the cystic nature and oviductal origin of lesions from tdTomato oviductal tissue.

Cycle evaluation: We performed an H&E histological cycle evaluation of the endometrium in WT mice after cloprostenol induction and following estrogen and progesterone stimulation on day 12 to confirm that recipient mice were in the secretory phase. 3 cloprostanol induction and 1 E/P induction mice were euthanized daily beginning 1 day prior to planned tissue harvest and for 4 additional days for evaluation of the uterine horns.

### Uterine secretome (secretory phase) mouse model

Following development of the tdT mouse model, we next sought to evaluate whether tissues other than endometrium can implant ectopically in the model and whether uterine secretome factors may play a role in initial implantation of ectopic tissues. Ectopic tdT oviductal tissue was injected ± synchronized post-ovulatory WT endometrium to provide secretome factors. The first experiment based on prior models, included mouse specific steps including pseudodecidualization by vaginal sesame oil administration in donors to increase the amount of endometrial tissue [21], and administration of SC estrogen 24 hours after tissue injection in recipients based on “nidal estrogen” usage in mouse pregnancy models after ovulation to assist implantation [25–27], as well as tissue injections 2 weeks apart to maximize lesion development. Because numerous ectopic lesions were implanted, the next experiment more closely simulated human secretory cycles by reduced tissue injections to one, eliminated pseudodecidualization, and compared the use of WT endometrium ± post-injection estrogen administration. We also tested flushed oviductal cells to evaluate possible oviductal stromal effect. Subsequent experiments were performed with estrogen and progesterone synchronization only in donor mice to provide adequate diestrus endometrial tissue, since lesions developed in both arms. Live and post-euthanasia imaging was performed in these initial experiments at 14 and 28 days, and then monthly for the oviductal cell experiment, but was deemed unnecessary for subsequent experiments once optimal timing of gross lesion evaluation was determined to be 28 days.

### Progesterone Knockout tdT model

To demonstrate the importance of the uterine secretome on ectopic lesions development we superimposed the progesterone-induced uterine gland knockout (PUGKO) model [28] onto our tdT mouse model of ectopic lesions. The PUGKO mouse model results in the ablation of uterine glands and thereby a reduction in uterine secretome factors including LIF [28, 29]. tdT mouse pups on postnatal days 2-10 with progesterone 50 ug/g body weight (bw) (5 mg/ml sesame oil) SC with a 26 G needle. The progesterone effect on the endometrium was evaluated by immunohistochemical staining for uterine gland marker, forkhead box A2 (FOXA2) [28]. Primary antibody recombinant anti-Fox2a antibody (ab108422) and secondary goat anti-rabbit IgG H&L HRP conjugated (ab6721) were obtained from abcam and used according to manufacturer’s instructions. DAB Substrate kit (abcam) was used according to manufacturer’s instructions for visualization, hematoxylin was used to counterstain nuclei as we previously described [30].

Twelve recipient mice were used in each arm. Normal and PUGKO tdT mice were used as donors. Experimental arms included minced tdT oviduct, minced PUGKO tdT oviduct ± WT endometrium, minced tdT endometrium ± WT endometrium (Fig 3).

**Fig 3.**
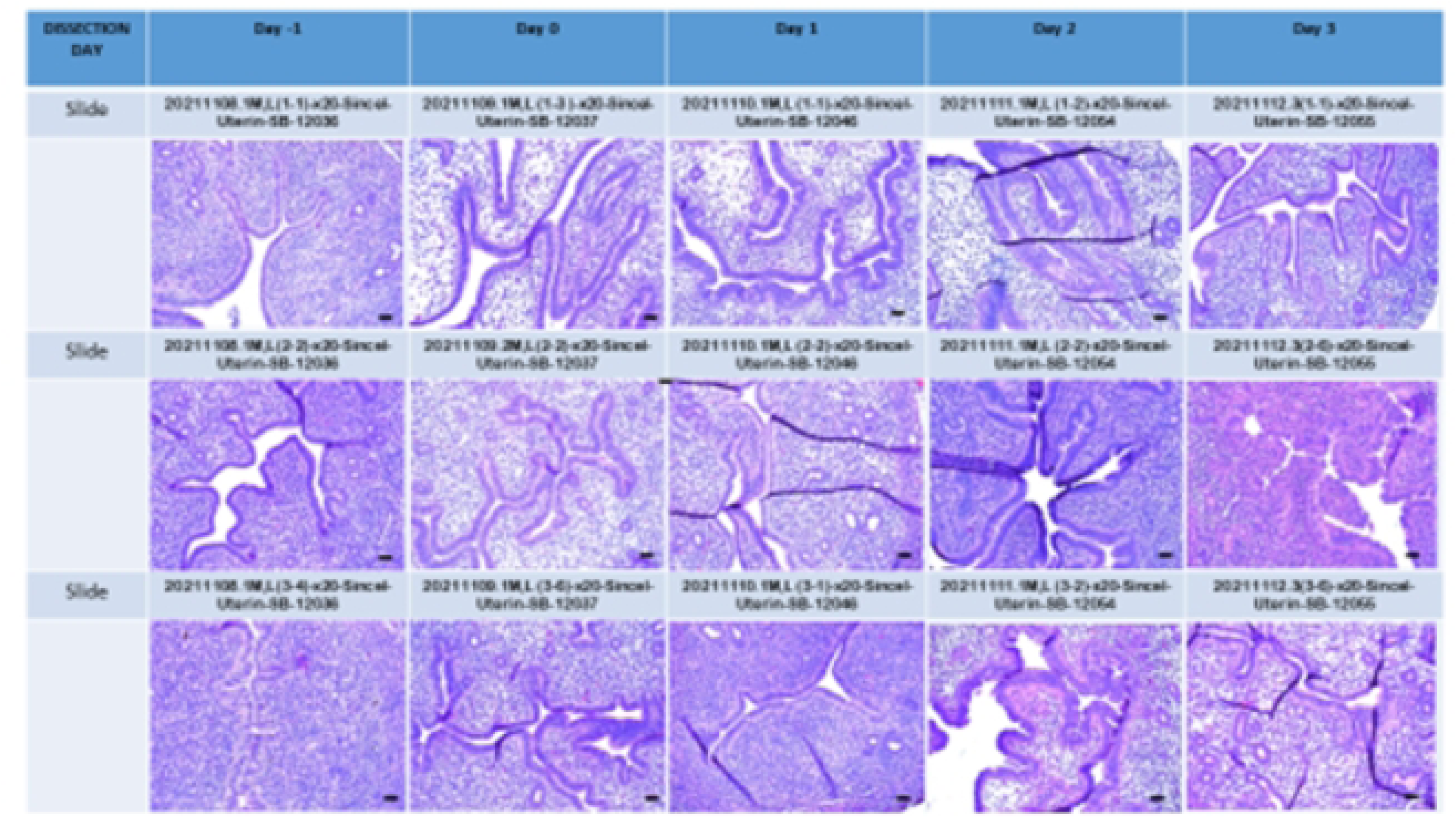
Summary slide of mouse uterine horn images from day –1 to 3 SINCEL treated method in x20 magnification. _50um. The order of injections; Day –5: Progesterone (3 µg) in 200µl saline injected SQ. Cloprostenol (0.5 µg) in 100µl saline injected separately IP. Day –2: Cloprostenol (0.5µg) in 100µl saline, injected IP.

### Hormonal effects

Next, we evaluated the effects of hormonal cycling on ectopic lesions development. Cyclicity was assessed in the recipient and donor mice via H&E histology. Synchronized female recipient mice were changed to male mice suppressed with degarelix acetate 0.5 mg SC (5mg/mL sterile water), a GnRH antagonist, 2-weeks before tissue injection to eliminate possible recipient hormonal or other contributions on lesion development, and to decrease mouse wastage. Female donor mice suppressed with degarelix were evaluated to assess the importance of cycling endometrium.

Tissues from synchronized degarelix treated tdT mice and/or endometrial tissue from synchronized WT mice were injected into the peritoneal cavity of degarelix treated male C57BL/6 WT recipient mice, using minced endometrial tissue alone, minced oviductal tissue alone, or a combination of oviductal tissue and minced endometrial tissue from donor tissue incubated with LIF as described below to evaluate the implantation of lesions in both arms.

### Leukemia inhibitory factor supplementation

Finally, the addition of recombinant LIF as a single experimental variable was tested to see if ectopic endosalpingeal and endometrial lesion development increases in its presence. Oviductal and endometrial epithelium was harvested and injected in a similar fashion ± LIF, using different incubation times and concentration. LIF 10 ug [100 ug/mL] in 100uL was added to 300uL PBS with tissue, incubated for 5 minutes and then injected into recipient mice in the experimental arms. Endometrial tissue with LIF 40 ug in 400 uL PBS with a 30 minutes incubation period was also evaluated.

### Prepubertal Ovarian inclusion cyst (OIC) development

We also evaluated the presence of OICs in pre-pubertal mice to assess whether spontaneous metaplasia might be a possible source of serous lesions. Mice are known to begin ovulating by approximately 35 days, so 100 ovaries of 50 WT C57BL/6 mice euthanized at 30-31 days were assessed for OICs. This number of mice were utilized, since OICs were not expected to be present. The ovaries were fixed in 10% buffered formalin at room temperature for 24 hours. The tissues were processed through ethanol dilutions into paraffin wax. Two transverse 5 µm sections were serially performed in every 20 µm intervals, one slide for H&E staining per standard protocols and the second adjacent slide preserved for IHC staining in case of OIC presence.

## Statistical Analysis

Statistical analysis was performed using GraphPad Prism (La Jolla, CA, USA). IPITT were analyzed using 2-way ANOVA with diet and time as factors. For multiple comparisons an ANOVA was used, and Tukey test was used for post-hoc, pairwise comparisons.

## Results

### Model establishment using tdTomato mice

To visualize ectopic lesion growth over time in-vivo, we developed a mouse model using tdT mice as donors and WT mice as recipients. Lesions were visualized 28 days post injection of either endometrial or oviductal donor tissue (Fig. 1B). The use of tdT tomato mice as donors also allowed for visualization of lesions at time of necropsy (Fig. 1C). All the mice in the initial experiment with two study arms (Fig. 1A), one arm receiving minced tdT oviduct alone, and the second receiving minced tdT oviduct and menstrual WT endometrium to provide secretome factors, had extensive implanted lesions, up to 14/mouse. Mice receiving both endometrium and oviduct averaged numerically but not significantly more lesions than those who received oviduct alone (5.8 EO lesions per mouse versus 4.4 O, p=0.489). Endosalpingiosis/oviductal tissue was implanted in all recipients in both study arms. Overall, the use of tdT mice as donors into WT recipient mice provides in-vivo visualization of lesion development and allows for visualization of small lesions at time of dissection.

### Uterine secretome (secretory phase) factors increase ectopic lesion development

Given that oviductal implantation in the model was successful, and occurred in the absence of supplemental WT endometrium, we proceeded with a series of experiments (S1 Table) to determine if the presence of uterine secretome factors play a role in the process of ectopic tissue growth, by eliminating possible confounding variables. Initial factors we considered were: an effect of oviductal stromal cells in the minced oviductal tissue, and two factors distinct in the mouse model compared to humans: uterine horn pseudodecidualization, and the effect of post-injection supplemental estrogen. Initially, all mice were sacrificed on days 21-28 after initial implantation. However, on initial imaging in the experiment using oviductal cells, no lesions were visualized. Due to the concern that individual cells would potentially require a longer time to grow into visible lesions, mice in this experiment were sacrificed 93 days after implantation. The oviductal cell experiment showed lesions in 3/10 mice, 2 in the post injection estrogen arm and 1 in the no estrogen arm (5 with estrogen and 5 without estrogen). Additional experiments were completed using minced tissues due to difficulty harvesting cells alone and the time duration for lesion growth, since our focus was factors capable of affecting implantation of free-floating tissues. Since lesion development occurred ± pseudodecidualization and also ± post injection estrogen treatment, these treatments were sequentially removed from the experiments to reduce confounders.

Minced tdT oviduct was then implanted ± WT endometrium, with 12 mice/arm (Fig 2B) There was significant increase in the number of oviductal lesions when implanted with endometrium versus without.

An experiment to confirm the effect of cloprostenol treatment to achieve the diestrus phase was performed by H&E evaluation of the endometrium following cloprostanol induction (Fig 3). The histologic evaluation confirmed that mice had secretory endometrium 48 hours after 2 cloprostenol injections (at day 2 following the second cloprostanol injection).

### Ablation of recipient uterine glands significantly decreases implantation of ectopic lesions

Subsequently, we sought to manipulate the uterine secretome and therefore we superimposed the PUGKO uterine receptivity mouse model onto our mouse model of ectopic lesions (Fig. 4. A&B). Uterine gland marker, FOXA2, was reduced in PUGKO compared to control mice (Fig. 4 C&D). Endometrial lesion count was significantly decreased in PUGKO tdT mice compared to tdT mice (p<0.05) (Fig 4E). Lesion development increased with addition of WT endometrium as expected, but this was not statistically significant with the sample size and small lesion numbers (12 lesions/6 mice vs 4 lesions/6 mice p=0.12) (Fig 4F). Oviductal lesion count was minimally decreased in the tdT PUGKO model compared to tdT tissue, as expected, since FOXA2 is thought to affect uterine secretome factors (Fig. 4G).

**Fig 4.**
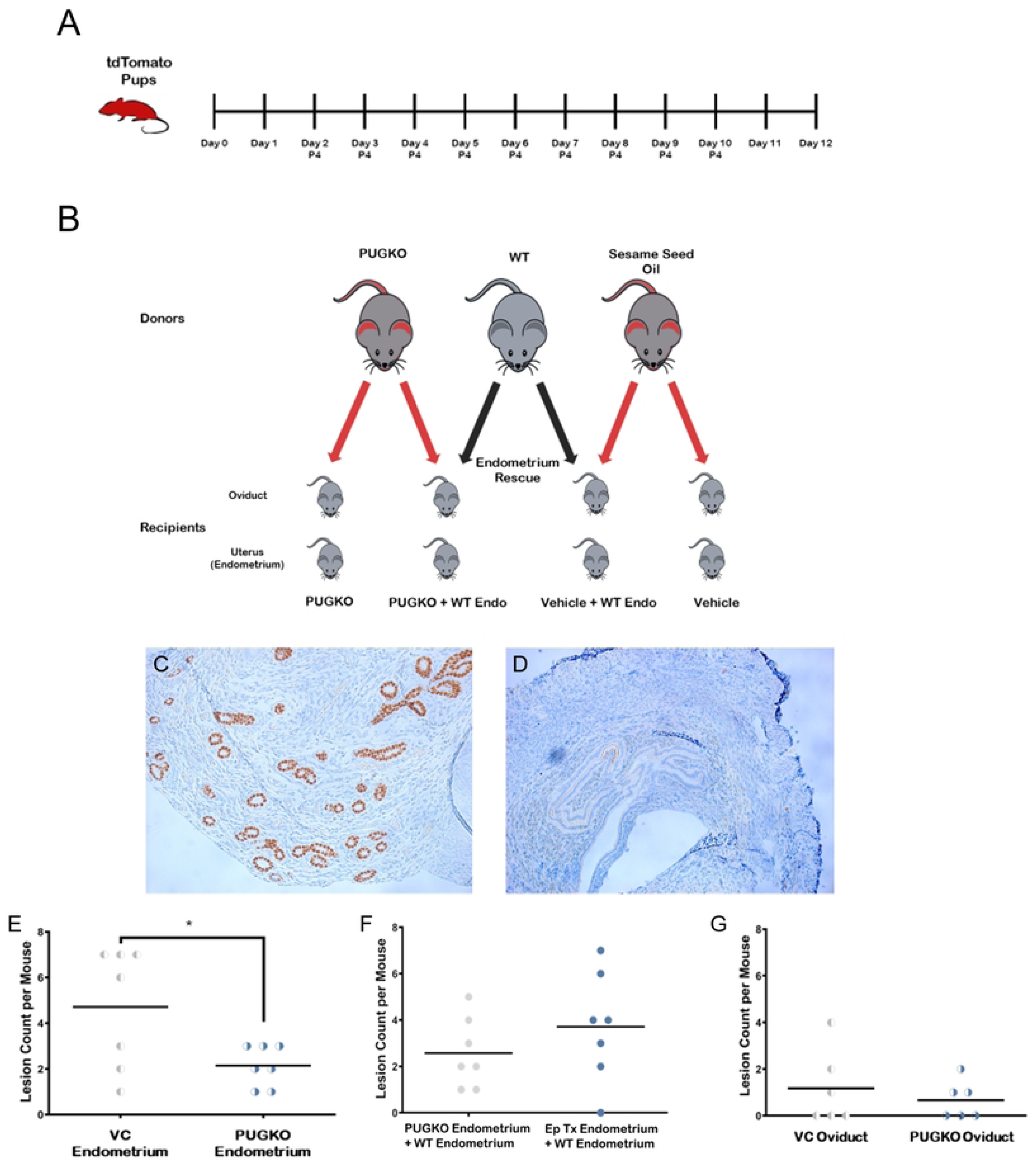
Ectopic lesions are decreased in PUGKO mice compared to controls. A. Creation of tdT PUGKO mouse model with daily injections of progesterone from postnatal day 2-10 to knock down secretome factors was based on mouse model that results in infertility in female mice. P4= progesterone B. Oviduct tissue and endometrium from progesterone treated (PUGKO) and vehicle control donor mice were collected and processed, then injected into WT C57Bl/6J recipient mice. Due to the fluorescent nature of the tdT mouse tissue, all tissue that implanted in the recipient mice could be identified using a red filter with a green light exposure. C –D. Immunohistochemistry (IHC) staining for FOXA2 was performed to assess the response of pups to progesterone treatment (PG TX). This was measured by comparing the counted IHC FOXA2 stained uterine glands of microscopic images in 4 high power fields (HPF) of 40 magnification in PUGKO tdT (D) vs WT (C). PG treatment in PUGKO mice decreases the expression of FOXA2 almost 33 times compared to the non-PG TX mice. **E.** Lesion count in mice with endometrial tissue from tdT mice ± progesterone on postnatal day 2-10 (PUGKO). There are significantly less lesions when mice recieve progesterone as pups (*=p<0.05). **F.** Lesion development using PUGKO endometrium and secretome factors (provided by synchronized WT endometrium). PUGKO reduces implantation factors and associates with fewer endometrial lesion growths. WT endometrium appears to restore implantation of PUGKO endometrium. There is no statistically significant difference when PUGKO endometrium is co-injected with WT endometrium. **G.** PUGKO does not significantly alter the number of oviductal lesions. The progesterone induced blocking of uterine glands in PUGKO mice did not statistically affect the number of oviductal lesions compared to controls in vehicle mice, since oviduct does not produce secretome factors. (Student t-test.) E: Estrogen, P: Progesterone

### Hormonal cycling in both donor and recipient mice affect lesion initiation

Our initial experiments demonstrated that oviductal tissue from tdT mice have increased lesion development in the presence of secretory factors provided by WT endometrium (Fig. 2B). In these experiments we wanted to explore the role recipient reproductive hormones, specifically estrogen and progesterone, affect ectopic lesion development. To do this, we utilized male mice treated with degarelix as recipients. Finally, absence of hormonal stimulation of donor tissue decreased lesion development when donor mice were treated with degarelix to suppress endometrial cycling. Of note, a low baseline implantation rate was noted for tissues that were hormonally suppressed (Fig. 5).

**Fig 5.**
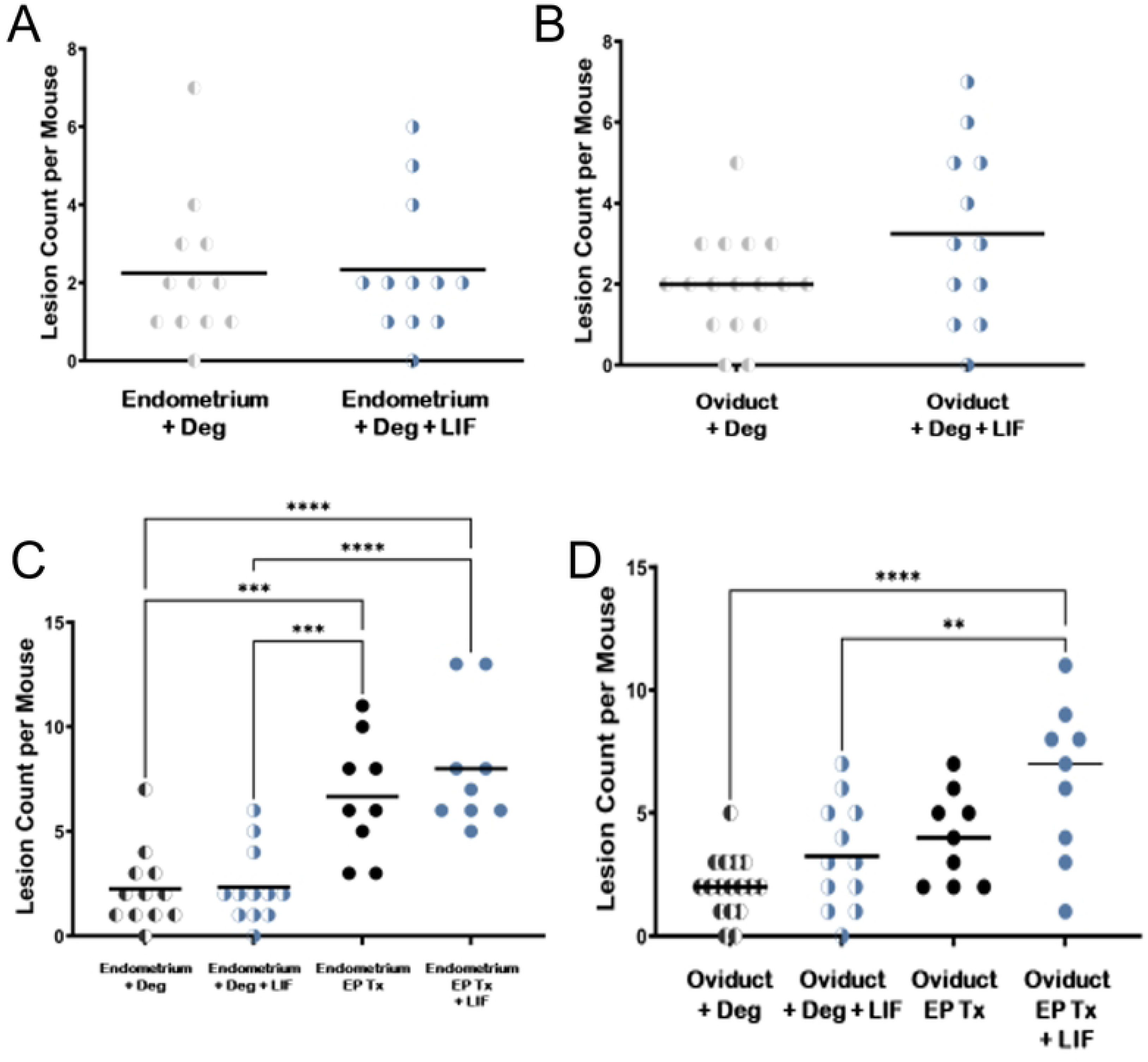
Effects of LIF on ectopic lesion growth **A.** The addition of LIF does not alter the number of endometrial lesions. When the donor and recipient male mice were hormonally suppressed with degarelix, the co-injection of LIF did not demonstrate any statistically significant change in endometrial lesion/mouse compared to solely implanted endometrial controls. **B.** Addition of LIF alone does not statistically alter the number of oviductal lesions. In hormonally suppressed donor and recipient male mice with degarelix, the co-injection of LIF did not demonstrate significant change in oviductal lesion/mouse compared to solely implanted oviductal controls treated with degarelix. C. Disruption of the estrus cycle by degarelix injection in donor mice significantly reduces the number of endometrial lesions/mouse in comparison with hormonally synchronized controls injected with endometrium (***p<0.001). Addition of exogenous LIF to hormonally synchronized endometrium further significantly increases the number of injected ectopic endometrial lesions/mouse (****P<0.0001). **D** Implantation of oviductal tissue from hormonally synchronized mice with LIF significantly increases the number of lesions/mouse compared to oviductal lesion number in degarelix treated mice ± LIF (**p<0.01) which are similar in number of lesions to hormonally synchronized mice without LIF. E:estrogen, P: progesterone

### Leukemia inhibitory factor improves lesion implantation of oviductal tissue

The addition of leukemia inhibitory factor (LIF), a cytokine factor long known to play a role in mouse pregnancy implantation, to endometrial tissue did not statistically increase the lesion count when given to degarelix suppressed mice with small numbers of mice. (Fig 5A&B). On the other hand, hormonally treated mice had significantly more lesions than mice injected with endometrium from mice treated with degarelix, with most lesions in the hormones + LIF treatment cohort, demonstrating that hormones are important to prepare the uterine secretome factors that play a role in lesion implantation, and LIF works in combination with the secretome factors (Fig 5C). Oviduct tissue treated with LIF before injection into donor mice also resulted in a significant increase in lesion count (Fig. 5D). The addition of the implantation factor LIF increased the implantation of oviductal tissue but not endometrial tissue, presumably because the endometrium secretes endogenous LIF.

### Prepubertal ovarian inclusion cyst assessment

Retrograde flow and implantation as the source of ectopic lesions has been an accepted theory for nearly 100 years and is explained by the secretome theory. An alternate theory is mataplasia of ovarian tissue, so we searched for the presence of OICs in “pre-ovulatory” mice, and found no lesions in these mice that have not yet initiated ovulation (Fig. 6)

**Fig 6.**
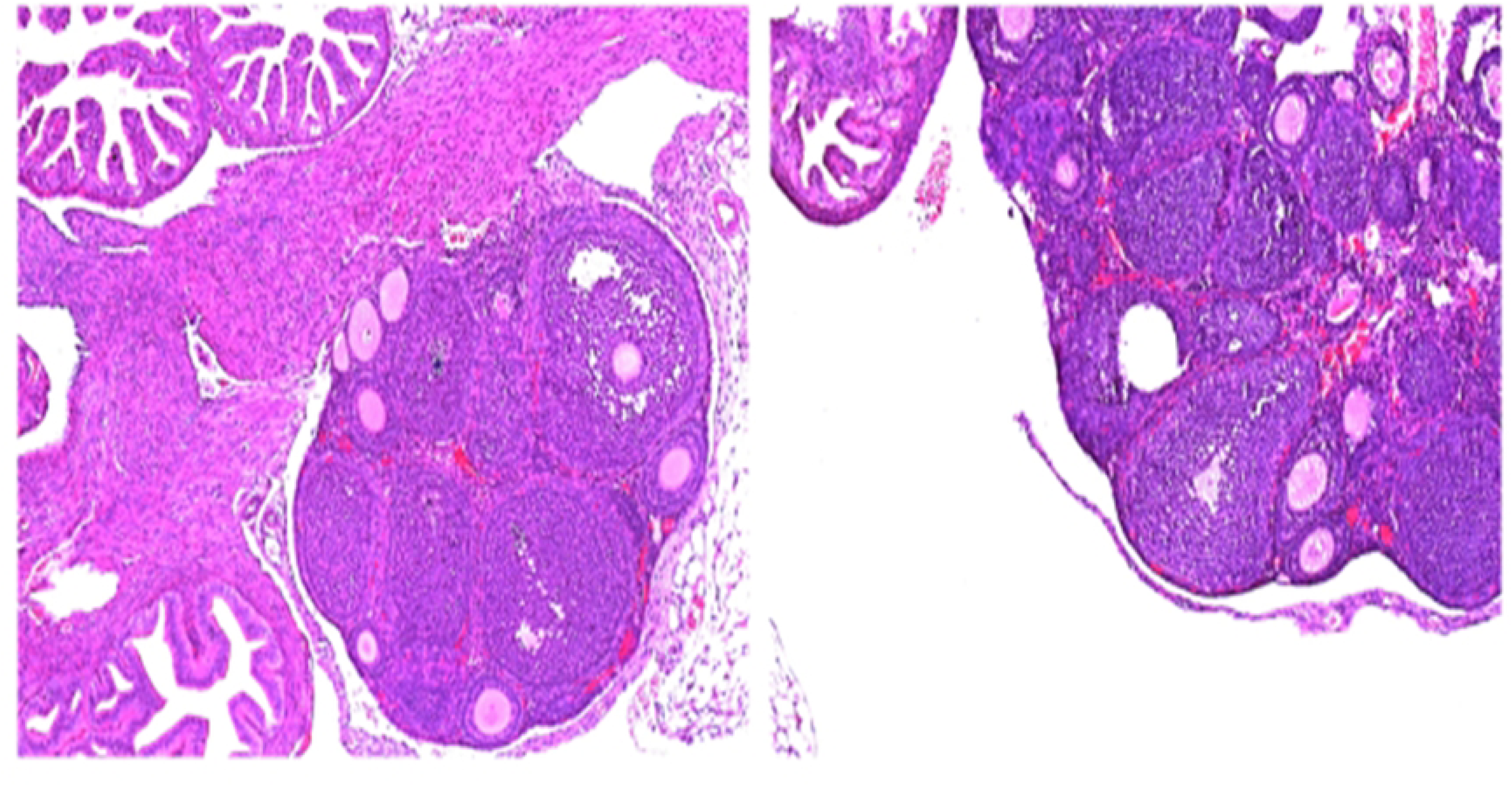
H&E stained 5 µm sections of two WT C57BL/6 mice ovaries at x10 demonstrate follicles, but no OICs.

## DISCUSSION

Our mouse model introduces the “uterine secretome” theory that free floating gynecologic epithelial cells which migrate, most often to the peritoneal cavity, are influenced by uterine secretome factors to implant ectopically. Neither Sampson’s retrograde menstruation theory [31], nor the “precursor escape” theory leading to serous ovarian cancer, first proposed in 2003 by Piek et al [32] and others [33–35] addresses how ectopic tissues initiate the attachment process, nor do they explain the reported 89% prevalence of benign epithelial ectopic lesions in menopausal patients [2]. The uterine secretome theory expands upon the “precursor escape” theory by explaining how benign and malignant lesions are initiated, and also explains the increase of ectopic growths with age, as more secretory cycles lead to more lesions. Distant ectopic growths are likely due to detached endometrial cells, which have been demonstrated in the bloodstream [36–38] driven by the secretome to implant.

Our tdTomato ectopic lesion model described here, which is based on earlier endometriosis models [21–23], successfully implanted many endosalpingial lesions in both arms initially, using estrogen and progesterone followed by pseudodecidualization to create a menstrual endometrium, not natural in mice, prior to tissue harvesting, and estrogen supplementation in the recipients ± post-ovulatory WT endometrial tissue to provide secretome factors. Subsequent iterations tested confounding variables in both the donor and recipient mice, beginning with mouse-specific factors. The first variables eliminated were mouse –specific pseudodecidualization associated with uterine stimulation, and the estrogen “nidatory surge” which did not prevent ectopic lesion development. Minced tissue in the diestrus or “secretory” phase of the cycle achieved by hormonal manipulation was sufficient to increase lesion development, and was easier to work with compared to flushed oviductal cells, so minced tissues were further used to test the effect of the uterine secretome on lesion numbers.

The PUGKO tdT mouse model treated mice with postnatal progesterone on days 2-10 (based on a fertility model that shows decreased embryo implantation as adults) to decrease FOXA2, an upstream regulator of LIF [28, 29], which was confirmed by IHC staining of the uterine horn. There is some variability in LIF effect in different mouse species [39] so we did not anticipate elimination of FOXA2. Significantly decreased endometriosis lesion development was found and was restored by supplementation of implanted PUGKO tdT tissues with WT endometrium to provide secretome factors, indicating that uterine factors increase ectopic tissue implantation. Oviductal lesion numbers in PUGKO tdT mice were not significantly different than tdT mice, since the oviduct does not produce secretome factors, but were increased in the presence of WT endometrium (Fig 7).

We demonstrated that there is a baseline level of lesion development even when the donor endometrium is suppressed and that hormonal manipulation to place the endometrium in the secretory phase significantly increases lesion numbers, with a further increase with addition of synchronized WT endometrium to provide secretome factors. Recipient female mice were initially synchronized to the diestrus/secretory phase using cloprostanol induction, and subsequently hormonally suppressed male mice were successfully utilized to eliminate the possibility of recipient uterine secretome factors or other hormonal effects.

To demonstrate the role of a known uterine secretome factor, we supplemented tissue injections with recombinant LIF, which statistically increased oviduct and endometrium implantation in the presence of hormonally synchronized endometrium. Gene expression studies in embryo implantation [28, 29, 40], endometriosis [10, 11] and ovarian cancer metastasis [41] have all found altered expression of LIF, and other genes such as CD44 and miR 99a-5p [42, 43], supporting our hypothesis that the uterine secretome is important in benign and malignant processes. Similar miRNA has been reported in these settings [44, 45]. LIF affects the JAK-STAT pathway, which is altered in ovarian metastasis [46]. Endometriosis has been shown to have elevated LIF in the peritoneal fluid [9, 10]. An earlier paper reports a 30-40% difference in LIF from uterine flushings of EM patients, which they considered non-significant [47], but the non-significance may be due to small numbers in their study. Additional recent confirmatory data supporting the secretome theory include the association of endometriosis with exosomes that increase invasion and migration [9, 47, 48], and the persistence of LIF into the menstrual phase in humans [49]. Further study of the overlap of altered gene expression in these processes will be a focus to elucidate the role of these genes and others in the ectopic benign and malignant lesion initiation and metastatic processes.

The “uterine secretome” and “precursor escape” theories are strongly supported by recent data. A 2022 large multi-institutional study showed that risk of primary peritoneal serous carcinoma (PPSC) is increased in women with serous tubal intraepithelial carcinomas (STIC) compared to those with normal Fallopian tube epithelium in women having risk reducing salpingo-oophorectomy (RRSO), with an estimated hazard ratio to develop PPSC during follow-up in women with STIC of 33.9 [50] likely due to implantation of peritoneal pre-malignant lesions prior to RRSO that become malignant over time. Additional studies that support this concept provide data that earlier RRSO provides greater protection [50, 51] and that opportunistic salpingectomy decreases multiple ovarian epithelial cancer subtypes [48, 52].

It was demonstrated in 1982 that epithelial ovarian cysts typically develop after puberty and that hormonal stimulation is critical to the process [53], likely via the secretome. Our mouse data is consistent with the finding that ectopic gynecologic tissue growth is hormonally driven, as we found no prepubertal OIC lesions in mice.

In conclusion, we developed a mouse model of ectopic lesions that incorporates the ability to visualize ectopic lesions growth in-vivo over time using tdT transgenic mice as donors. Furthermore, the data presented here demonstrates that uterine derived secretome factor/s are the critical factor/s in initiation of ectopic lesion growth. We also showed that LIF, an essential uterine secretome factor, stimulates ectopic lesion growth. Further studies will utilize our new mouse model to further our understanding of how ectopic lesion growth is initiated in both benign and malignant ectopic lesions.

## REFERENCES

1. Esselen KM, Terry KL, Samuel A, Elias KM, Davis M, Welch WR, et al. Endosalpingiosis: More than just an incidental finding at the time of gynecologic surgery? Gynecol Oncol. 2016;142(2):255–60. doi: 10.1016/j.ygyno.2016.05.036. PubMed PMID: 27261327.

2. Sunde J, Wasickanin M, Katz TA, Wickersham EL, Steed DOE, Simper N. Prevalence of endosalpingiosis and other benign gynecologic lesions. PloS one. 2020;15(5):e0232487. doi: 10.1371/journal.pone.0232487. PubMed PMID: 32401810; PubMed Central PMCID: PMCPMC7219775 members of the armed forces. The views expressed are those of the author(s) and do not reflect the official policy of the Department of the Army, the Department of Defense of the US government. The investigators have adhered to the policies for protection of human subjects as prescribed in 45 CFR 46.

3. Gubbels AL, Li R, Kreher D, Mehandru N, Castellanos M, Desai NA, et al. Prevalence of occult microscopic endometriosis in clinically negative peritoneum during laparoscopy for chronic pelvic pain. Int J Gynaecol Obstet. 2020;151(2):260–6. doi: 10.1002/ijgo.13303. PubMed PMID: 32644227.

4. Seidman JD, Khedmati F. Exploring the histogenesis of ovarian mucinous and transitional cell (Brenner) neoplasms and their relationship with Walthard cell nests: a study of 120 tumors. Arch Pathol Lab Med. 2008;132(11):1753–60. doi: 10.5858/132.11.1753. PubMed PMID: 18976011.

5. Roma AA, Masand RP. Ovarian Brenner tumors and Walthard nests: a histologic and immunohistochemical study. Hum Pathol. 2014;45(12):2417–22. doi: 10.1016/j.humpath.2014.08.003. PubMed PMID: 25281026.

6. Paner GP, Gonzalez M, Al-Masri H, Smith DM, Husain AN. Parafallopian tube transitional cell carcinoma. Gynecol Oncol. 2002;86(3):379–83. doi: 10.1006/gyno.2002.6777. PubMed PMID: 12217766.

7. Lagana AS, Vitale SG, Salmeri FM, Triolo O, Ban Frangez H, Vrtacnik-Bokal E, et al. Unus pro omnibus, omnes pro uno: A novel, evidence-based, unifying theory for the pathogenesis of endometriosis. Med Hypotheses. 2017;103:10–20. doi: 10.1016/j.mehy.2017.03.032. PubMed PMID: 28571791.

8. Zhou J, Chern BSM, Barton-Smith P, Phoon JWL, Tan TY, Viardot-Foucault V, et al. Peritoneal Fluid Cytokines Reveal New Insights of Endometriosis Subphenotypes. Int J Mol Sci. 2020;21(10). doi: 10.3390/ijms21103515. PubMed PMID: 32429215; PubMed Central PMCID: PMCPMC7278942.

9. Jorgensen H, Hill AS, Beste MT, Kumar MP, Chiswick E, Fedorcsak P, et al. Peritoneal fluid cytokines related to endometriosis in patients evaluated for infertility. Fertil Steril. 2017;107(5):1191–9 e2. doi: 10.1016/j.fertnstert.2017.03.013. PubMed PMID: 28433374.

10. Li C, Zhao HL, Li YJ, Zhang YY, Liu HY, Feng FZ, et al. The expression and significance of leukemia inhibitory factor, interleukin-6 and vascular endothelial growth factor in Chinese patients with endometriosis. Arch Gynecol Obstet. 2021;304(1):163–70. doi: 10.1007/s00404-021-05980-5. PubMed PMID: 33555431.

11. Zutautas KB, Sisnett DJ, Miller JE, Lingegowda H, Childs T, Bougie O, et al. The dysregulation of leukemia inhibitory factor and its implications for endometriosis pathophysiology. Front Immunol. 2023;14:1089098. doi: 10.3389/fimmu.2023.1089098. PubMed PMID: 37033980; PubMed Central PMCID: PMCPMC10076726.

12. Stewart CL, Kaspar P, Brunet LJ, Bhatt H, Gadi I, Kontgen F, et al. Blastocyst implantation depends on maternal expression of leukaemia inhibitory factor. Nature. 1992;359(6390):76-9. doi: 10.1038/359076a0. PubMed PMID: 1522892.

13. Booth M, Beral V, Smith P. Risk factors for ovarian cancer: a case-control study. Br J Cancer. 1989;60(4):592–8. doi: 10.1038/bjc.1989.320. PubMed PMID: 2679848; PubMed Central PMCID: PMCPMC2247100.

14. Wynder EL, Dodo H, Barber HR. Epidemiology of cancer of the ovary. Cancer. 1969;23(2):352–70. doi: 10.1002/1097-0142(196902)23:2<352::aid-cncr2820230212>3.0.co;2-4. PubMed PMID: 5764976.

15. Gaitskell K, Coffey K, Green J, Pirie K, Reeves GK, Ahmed AA, et al. Tubal ligation and incidence of 26 site-specific cancers in the Million Women Study. Br J Cancer. 2016;114(9):1033–7. doi: 10.1038/bjc.2016.80. PubMed PMID: 27115569; PubMed Central PMCID: PMCPMC4984917.

16. Gaitskell K, Green J, Pirie K, Reeves G, Beral V, Million Women Study C. Tubal ligation and ovarian cancer risk in a large cohort: Substantial variation by histological type. Int J Cancer. 2016;138(5):1076–84. doi: 10.1002/ijc.29856. PubMed PMID: 26378908; PubMed Central PMCID: PMCPMC4832307.

17. Tan OL, Hurst PR, Fleming JS. Location of inclusion cysts in mouse ovaries in relation to age, pregnancy, and total ovulation number: implications for ovarian cancer? J Pathol. 2005;205(4):483–90. doi: 10.1002/path.1719. PubMed PMID: 15685692.

18. McMullen ML, Cho BN, Yates CJ, Mayo KE. Gonadal pathologies in transgenic mice expressing the rat inhibin alpha-subunit. Endocrinology. 2001;142(11):5005–14. doi: 10.1210/endo.142.11.8472. PubMed PMID: 11606469.

19. Bristol-Gould SK, Hutten CG, Sturgis C, Kilen SM, Mayo KE, Woodruff TK. The development of a mouse model of ovarian endosalpingiosis. Endocrinology. 2005;146(12):5228–36. doi: 10.1210/en.2005-0697. PubMed PMID: 16141389.

20. Burns KA, Pearson AM, Slack JL, Por ED, Scribner AN, Eti NA, et al. Endometriosis in the Mouse: Challenges and Progress Toward a ‘Best Fit’ Murine Model. Front Physiol. 2021;12:806574. doi: 10.3389/fphys.2021.806574. PubMed PMID: 35095566; PubMed Central PMCID: PMCPMC8794744.

21. Greaves E, Cousins FL, Murray A, Esnal-Zufiaurre A, Fassbender A, Horne AW, et al. A novel mouse model of endometriosis mimics human phenotype and reveals insights into the inflammatory contribution of shed endometrium. Am J Pathol. 2014;184(7):1930–9. doi: 10.1016/j.ajpath.2014.03.011. PubMed PMID: 24910298; PubMed Central PMCID: PMCPMC4076466.

22. Cousins FL, Murray A, Esnal A, Gibson DA, Critchley HO, Saunders PT. Evidence from a mouse model that epithelial cell migration and mesenchymal-epithelial transition contribute to rapid restoration of uterine tissue integrity during menstruation. PloS one. 2014;9(1):e86378. doi: 10.1371/journal.pone.0086378. PubMed PMID: 24466063; PubMed Central PMCID: PMCPMC3899239.

23. Dodds KN, Beckett EAH, Evans SF, Hutchinson MR. Lesion development is modulated by the natural estrous cycle and mouse strain in a minimally invasive model of endometriosis. Biol Reprod. 2017;97(6):810–21. doi: 10.1093/biolre/iox132. PubMed PMID: 29069288.

24. Pallares P, Gonzalez-Bulnes A. A new method for induction and synchronization of oestrus and fertile ovulations in mice by using exogenous hormones. Lab Anim. 2009;43(3):295–9. doi: 10.1258/la.2008.008056. PubMed PMID: 19116296.

25. Somigliana E, Vigano P, Zingrillo B, Ranieri S, Filardo P, Candiani M, et al. Induction of endometriosis in the mouse inhibits spleen leukocyte function. Acta obstetricia et gynecologica Scandinavica. 2001;80(3):200–5. doi: 10.1034/j.1600-0412.2001.080003200.x. PubMed PMID: 11207484.

26. Hoversland RC, Dey SK, Johnson DC. Catechol estradiol induced implantation in the mouse. Life Sci. 1982;30(21):1801–4. doi: 10.1016/0024-3205(82)90316-2. PubMed PMID: 6285111.

27. Shelesnyak MC. A history of research on nidation. Ann N Y Acad Sci. 1986;476:5–24. doi: 10.1111/j.1749-6632.1986.tb20918.x. PubMed PMID: 3541747.

28. Kelleher AM, Burns GW, Behura S, Wu G, Spencer TE. Uterine glands impact uterine receptivity, luminal fluid homeostasis and blastocyst implantation. Sci Rep. 2016;6:38078. doi: 10.1038/srep38078. PubMed PMID: 27905495; PubMed Central PMCID: PMCPMC5131473.

29. Kelleher AM, DeMayo FJ, Spencer TE. Uterine Glands: Developmental Biology and Functional Roles in Pregnancy. Endocr Rev. 2019;40(5):1424–45. doi: 10.1210/er.2018-00281. PubMed PMID: 31074826; PubMed Central PMCID: PMCPMC6749889.

30. Pennington KA, Oestreich AK, Cataldo KH, Fogliatti CM, Lightner C, Lydon JP, et al. Conditional knockout of leptin receptor in the female reproductive tract reduces fertility due to parturition defects in mice. Biol Reprod. 2022;107(2):546–56. doi: 10.1093/biolre/ioac062. PubMed PMID: 35349646.

31. Sampson JA. Metastatic or Embolic Endometriosis, due to the Menstrual Dissemination of Endometrial Tissue into the Venous Circulation. Am J Pathol. 1927;3(2):93–110 43. PubMed PMID: 19969738; PubMed Central PMCID: PMCPMC1931779.

32. Piek JM, Verheijen RH, Kenemans P, Massuger LF, Bulten H, van Diest PJ. BRCA1/2-related ovarian cancers are of tubal origin: a hypothesis. Gynecol Oncol. 2003;90(2):491. doi: 10.1016/s0090-8258(03)00365-2. PubMed PMID: 12893227.

33. Piek JM, Kenemans P, Verheijen RH. Intraperitoneal serous adenocarcinoma: a critical appraisal of three hypotheses on its cause. Am J Obstet Gynecol. 2004;191(3):718–32. doi: 10.1016/j.ajog.2004.02.067. PubMed PMID: 15467531.

34. Kurman RJ, Vang R, Junge J, Hannibal CG, Kjaer SK, Shih Ie M. Papillary tubal hyperplasia: the putative precursor of ovarian atypical proliferative (borderline) serous tumors, noninvasive implants, and endosalpingiosis. Am J Surg Pathol. 2011;35(11):1605–14. doi: 10.1097/PAS.0b013e318229449f. PubMed PMID: 21997682; PubMed Central PMCID: PMCPMC3193599.

35. Levanon K, Crum C, Drapkin R. New insights into the pathogenesis of serous ovarian cancer and its clinical impact. J Clin Oncol. 2008;26(32):5284–93. doi: 10.1200/JCO.2008.18.1107. PubMed PMID: 18854563; PubMed Central PMCID: PMCPMC2652087.

36. Bobek V, Kolostova K, Kucera E. Circulating endometrial cells in peripheral blood. Eur J Obstet Gynecol Reprod Biol. 2014;181:267–74. doi: 10.1016/j.ejogrb.2014.07.037. PubMed PMID: 25195200.

37. Grasso A, Navarro R, Balaguer N, Moreno I, Alama P, Jimenez J, et al. Endometrial Liquid Biopsy Provides a miRNA Roadmap of the Secretory Phase of the Human Endometrium. The Journal of clinical endocrinology and metabolism. 2020;105(3). doi: 10.1210/clinem/dgz146. PubMed PMID: 31665361.

38. Andronico F, Battaglia R, Ragusa M, Barbagallo D, Purrello M, Di Pietro C. Extracellular Vesicles in Human Oogenesis and Implantation. Int J Mol Sci. 2019;20(9). doi: 10.3390/ijms20092162. PubMed PMID: 31052401; PubMed Central PMCID: PMCPMC6539954.

39. Kobayashi R, Terakawa J, Kato Y, Azimi S, Inoue N, Ohmori Y, et al. The contribution of leukemia inhibitory factor (LIF) for embryo implantation differs among strains of mice. Immunobiology. 2014;219(7):512–21. doi: 10.1016/j.imbio.2014.03.011. PubMed PMID: 24698551.

40. Anegon I, Cuturi MC, Godard A, Moreau M, Terqui M, Martinat-Botte F, et al. Presence of leukaemia inhibitory factor and interleukin 6 in porcine uterine secretions prior to conceptus attachment. Cytokine. 1994;6(5):493–9. doi: 10.1016/1043-4666(94)90076-0. PubMed PMID: 7827286.

41. Chen LL, Ye F, Lu WG, Yu Y, Chen HZ, Xie X. Evaluation of immune inhibitory cytokine profiles in epithelial ovarian carcinoma. J Obstet Gynaecol Res. 2009;35(2):212–8. doi: 10.1111/j.1447-0756.2008.00935.x. PubMed PMID: 19335794.

42. Nakamura K, Sawada K, Kinose Y, Yoshimura A, Toda A, Nakatsuka E, et al. Exosomes Promote Ovarian Cancer Cell Invasion through Transfer of CD44 to Peritoneal Mesothelial Cells. Mol Cancer Res. 2017;15(1):78–92. doi: 10.1158/1541-7786.MCR-16-0191. PubMed PMID: 27758876.

43. Hua R, Wang Y, Lian W, Li W, Xi Y, Xue S, et al. Small RNA-seq analysis of extracellular vesicles from porcine uterine flushing fluids during peri-implantation. Gene. 2021;766:145117. doi: 10.1016/j.gene.2020.145117. PubMed PMID: 32920039.

44. Hu Q, Zang X, Ding Y, Gu T, Shi J, Li Z, et al. Porcine uterine luminal fluid-derived extracellular vesicles improve conceptus-endometrial interaction during implantation. Theriogenology. 2022;178:8–17. doi: 10.1016/j.theriogenology.2021.10.021. PubMed PMID: 34735978.

45. Mortlock S, Corona RI, Kho PF, Pharoah P, Seo JH, Freedman ML, et al. A multi-level investigation of the genetic relationship between endometriosis and ovarian cancer histotypes. Cell Rep Med. 2022;3(3):100542. doi: 10.1016/j.xcrm.2022.100542. PubMed PMID: 35492879; PubMed Central PMCID: PMCPMC9040176.

46. McLean K, Tan L, Bolland DE, Coffman LG, Peterson LF, Talpaz M, et al. Leukemia inhibitory factor functions in parallel with interleukin-6 to promote ovarian cancer growth. Oncogene. 2019;38(9):1576–84. doi: 10.1038/s41388-018-0523-6. PubMed PMID: 30305729; PubMed Central PMCID: PMCPMC6374186.

47. Mikolajczyk M, Wirstlein P, Skrzypczak J. Leukaemia inhibitory factor and interleukin 11 levels in uterine flushings of infertile patients with endometriosis. Hum Reprod. 2006;21(12):3054–8. doi: 10.1093/humrep/del225. PubMed PMID: 17000646.

48. Jiang Y, Chai X, Chen S, Chen Z, Tian H, Liu M, et al. Exosomes from the Uterine Cavity Mediate Immune Dysregulation via Inhibiting the JNK Signal Pathway in Endometriosis. Biomedicines. 2022;10(12). doi: 10.3390/biomedicines10123110. PubMed PMID: 36551866; PubMed Central PMCID: PMCPMC9775046.

49. Naseri S, Rosenberg-Hasson Y, Maecker HT, Avrutsky MI, Blumenthal PD. A cross-sectional study comparing the inflammatory profile of menstrual effluent vs. peripheral blood. Health Sci Rep. 2023;6(1):e1038. doi: 10.1002/hsr2.1038. PubMed PMID: 36620506; PubMed Central PMCID: PMCPMC9813904.

50. Steenbeek MP, van Bommel MHD, Bulten J, Hulsmann JA, Bogaerts J, Garcia C, et al. Risk of Peritoneal Carcinomatosis After Risk-Reducing Salpingo-Oophorectomy: A Systematic Review and Individual Patient Data Meta-Analysis. J Clin Oncol. 2022;40(17):1879–91. doi: 10.1200/JCO.21.02016. PubMed PMID: 35302882; PubMed Central PMCID: PMCPMC9851686 KrajcConsulting or Advisory Role: AstraZeneca, Pfizer Vilius RudaitisHonoraria: MedtronicConsulting or Advisory Role: AstraZeneca/MedImmuneResearch Funding: Inovio PharmaceuticalsTravel, Accommodations, Expenses: MSD Oncology, Roche, Karl Storz Elizabeth M. SwisherLeadership: IDEAYA BiosciencesNo other potential conflicts of interest were reported.

51. Harmsen MG, Piek JMJ, Bulten J, Casey MJ, Rebbeck TR, Mourits MJ, et al. Peritoneal carcinomatosis after risk-reducing surgery in BRCA1/2 mutation carriers. Cancer. 2018;124(5):952–9. doi: 10.1002/cncr.31211. PubMed PMID: 29315498.

52. Hanley GE, Pearce CL, Talhouk A, Kwon JS, Finlayson SJ, McAlpine JN, et al. Outcomes From Opportunistic Salpingectomy for Ovarian Cancer Prevention. JAMA Netw Open. 2022;5(2):e2147343. doi: 10.1001/jamanetworkopen.2021.47343. PubMed PMID: 35138400; PubMed Central PMCID: PMCPMC8829665 Zeneca outside the submitted work. No other disclosures were reported.

53. Blaustein A, Kantius M, Kaganowicz A, Pervez N, Wells J. Inclusions in ovaries of females aged day 1-30 years. Int J Gynecol Pathol. 1982;1(2):145–53. doi: 10.1097/00004347-198202000-00003. PubMed PMID: 7184893.

